# From multi-omics data to the cancer druggable gene discovery: a novel machine learning-based approach

**DOI:** 10.1101/2022.07.24.501277

**Authors:** Hai Yang, Lipeng Gan, Rui Chen, Dongdong Li, Jing Zhang, Zhe Wang

## Abstract

The development of targeted drugs allows precision medicine in cancer treatment and achieving optimal targeted therapies. Accurate identification of cancer drug genes is helpful to strengthen the understanding of targeted cancer therapy and promote precise cancer treatment. However, rare cancer-druggable genes have been found due to the multi-omics data’s diversity and complexity. This study proposes DF-CAGE, a novel machine learning-based method for cancer-druggable gene discovery. DF-CAGE integrated the somatic mutations, copy number variants, DNA methylation, and RNA-Seq data across ~10000 TCGA profiles to identify the landscape of the cancer-druggable genes. We found that DF-CAGE discovers the commonalities of currently known cancer-druggable genes from the perspective of multi-omics data and achieved excellent performance on OncoKB, Target, and Drugbank data sets. Among the ~20,000 protein-coding genes, DF-CAGE pinpointed 465 potential cancer-druggable genes. We found that the candidate cancer druggable genes (CDG-genes) are clinically meaningful and can be divided into highly reliable, reliable, and potential gene sets. Finally, we analyzed the contribution of the omics data to the identification of druggable genes. We found that DF-CAGE reports druggable genes mainly based on the CNAs data, the gene rearrangements, and the mutation rates in the population. These findings may enlighten the study and development of new drugs in the future.

## Introduction

Cancer is a genetic disease that leads the cause of global mortality [1, 2]. It originates in most cell types and organs of the body and is characterized by relatively unrestricted cell proliferation that can invade normal tissue boundaries and then metastasize to other organs [3]. Previous studies have shown that tumors arise because of the changes in the genome of cancer cells, including somatic mutations, copy number variations (CNVs), transcriptional profiles, epigenetic modifications, and altered DNA sequences [4]. The study of efficient and feasible cancer therapies has been a crucial issue [5]. The development of targeted drugs allows precision medicine in cancer treatment and achieving optimal targeted therapies [6]. The therapeutic advantages of targeted drugs are of great significance for improving the quality of life and prolonging the survival time of cancer patients [7].

Very few cancer druggable genes have been found so far [8]. The development of targeted drugs is mainly based on the cancer driver genes [9, 10]. The discovery of cancer driver genes is critical for cancer diagnosis, prevention, clinical treatment, and development of cancer-targeted drugs [11]. After years of development, many methods were proposed to pinpoint cancer driver genes, primarily based on mutation frequency evaluation. Lawrence et al. proposed Mustig2CV to identify significantly mutated by using gene expression data and DNA replication time data [12]. Mularoni et al. proposed OncodriverFML, which introduces a local mutation background model that calculates functional mutation bias genomic elements, analyzes somatic mutation patterns in coding and non-coding genomic regions and identifies driver mutations [13]. Tamborero et al. used silent mutations in the coding regions to construct a background model and proposed the OncodriverCLUST method, which mainly identifies genes with significant mutational clustering tendencies in protein sequences [14]. Reimand et al. proposed ActiveDriver, a Poisson generalized linear regression model-based approach to identify driver genes from gene sequences, identifying driver genes with high-frequency mutations after protein signal translational modifications [15]. With the increase in the number of driver genes, more and more machine learning approaches have become the focus of research, among which, Tokheim et al. proposed the 20/20+ method, a random forest-based machine learning prediction algorithm that uses features that capture mutation clustering, evolutionary conservation, prediction of variants, type of mutation outcome, gene interaction network connectivity, and other covariates for predicting oncogenes and tumor suppressor genes in somatic mutations [16]. Katchin et al. proposed CHASM, which uses sequence conservation and protein structure features to construct a random forest-based algorithm to predict driver mutations that are most likely to produce functional changes and promote cancer cell proliferation [17]. Han et al. proposed DriverML, which uses a statistical approach to quantify the functional impact scores of different invariant types on proteins and then combines machine learning models to identify cancer drivers [18].

With the rapid development and promotion of next-generation high-throughput sequencing technologies, international large-scale cancer projects such as The Cancer Genome Atlas (TCGA) [19], International Cancer Genome Collaboration (ICGC) [20], and Pan-Cancer Analysis of the Whole Genome (PCAWG) [21] have delivered a large number of multi-omics and clinical data based on different platforms with different technologies, providing a source of data to advance cancer study. Many early studies on cancer genes were analyzed using single dimension of omics data. However, due to the complexity of cancer, it is difficult to describe accurately by single dimension of omics data. Integrative analysis of multi-omics data can provide more insight into the flow of information in tumors [22, 23, 24].

Bailey et al. revealed that cancer driver genes account for more than 70% of the currently available druggable genes [25]. However, limited by the progress of driver gene discovery, the number of identified driver genes is rare, and only a tiny fraction of these genes is druggable. Thus, the current set of druggable cancer genes is still sparse [26, 27]. Finding more cancer druggable genes has become a bottleneck in cancer genomics study. This study proposes a machine learning-based approach with the deep forest for cancer druggable genes discovery (DF-CAGE) to overcome this bottleneck(the network framework of DF-CAGE is shown in Fig.1.). It integrates and extracts features from four types of omics data, namely DNA methylation, mRNA-seq data, CNVs, and somatic mutations. We found that DF-CAGE can directly identify cancer druggable genes using molecular omics data. The AUROC of DF-CAGE is ~0.9 on the OncoKB and TARGET datasets and is better than other state-of-the-art driver genes discovery approaches. We finally pinpointed 465 cancer druggable genes out of 20,000 proteincoding genes through model training and threshold selection. We classified these genes into three groups according to the different levels of reliability supported by the literature: 127 highly reliable genes (with existing drugs), 34 reliable genes (with clinical evidence), and 304 candidate genes (with DF-CAGE high probability scores). Finally, we analyzed the association of multi-omics data with the DF-CAGE identification results. We found that driver gene discovery methods mainly focus on mutation-related features in omics data. At the same time, DF-CAGE reports cancer druggable genes primarily based on the CNAs, gene rearrangements, and mutation rates in the population.

**Fig. 1.**
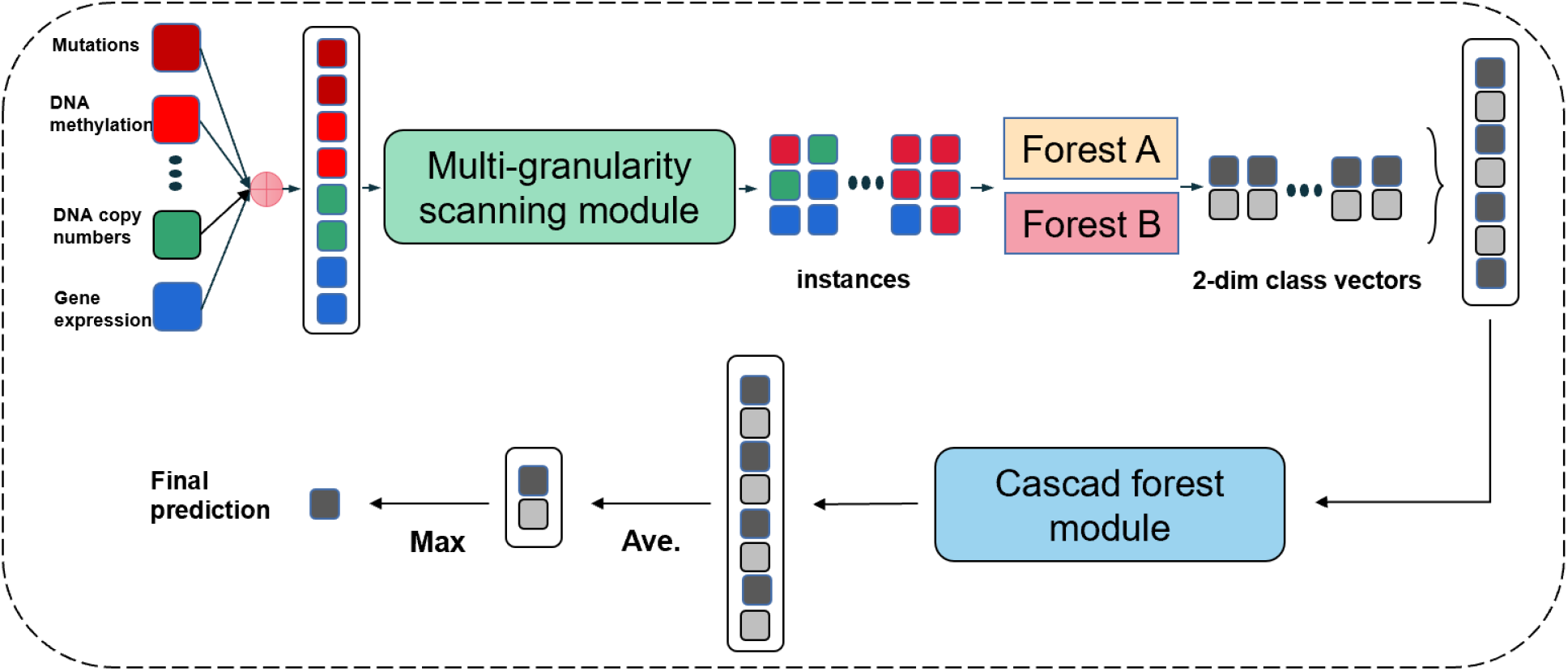
Overview of the DF-CAGE framework: the input module integrates multiple omics data; the multi-granularity scanning module extracts latent variables from the multi-omics data; the Cascading Forest Module generates each gene’s probability score.

## Material And Methods

The input of DF-CAGE is the multi-omics data of the TCGA pancancer dataset, and the output of DF-CAGE is the probability scores of the cancer druggable genes. The method can be divided into two steps. First, we extracted the features of DF-CAGE from the four omics data and applied DF-CAGE to explore the relationship between druggable genes and multi-omics data. Next, we used cascade forest to generate the prediction scores for cancer druggable genes.

### Data pre-processing and feature selection

This study’s cancer druggable genes discovery focused on the TCGA pan-cancer dataset (including 33 cancer types [28], as listed in Supplementary Table 1). We obtained DNA methylations, mRNA-seq data, copy number variations, and somatic mutations across the 33 cancers in the PanCancerAtlas. The four types of omics data were separately pre-processed and then integrated (refer to Supplementary Material 1 for detailed procedures of data preprocessing and feature selection.).

### Deep Forest model training

After the feature extraction process, DF-CAGE uses a deep forest approach based on an integrated learning method of multilayer forests to model the multi-omics features (refer to Supplementary Material 2 for DF-CAGE model parameter settings and training procedure.) The input to DF-CAGE is the feature matrix, which contains a 52-dimensional feature vector of all training set genes. CAGE model consists of two main components: the multi-granularity scan module and the cascade forest module. With the multi-granularity scanning module, DF-CAGE can learn the association between different omics data, enhancing the representation of the data and learning the differences and associations between multi-omics data more effectively in order to more accurately identify effective drug targets [29]. To improve the prediction accuracy of random forests on samples, DF-CAGE also uses a cascade forest module, which consists of multiple decision trees cascaded together. The cascade forest is similar to a multilayer perceptron. It uses sliding windows to extract feature vectors, which can enhance the nonlinear characterization ability of the model and maximize the relationship between the order of sample features and the prediction accuracy. The classification results are more accurate than traditional decision tree methods. The multi-granularity scanning process is shown in Fig.2.

**Fig. 2.**
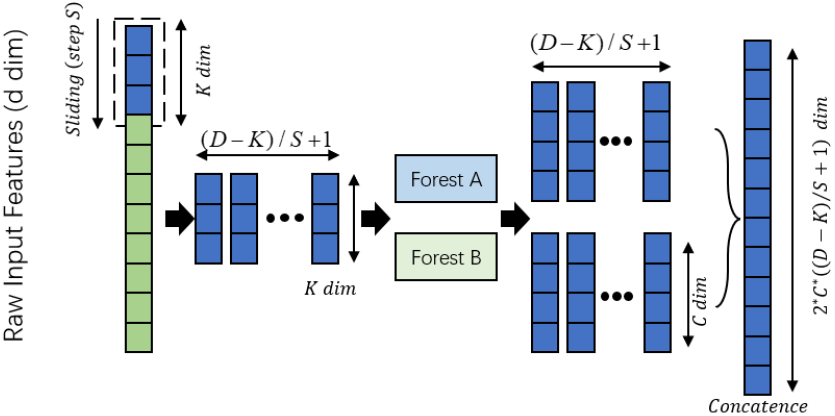
The multi-granularity scanning module can realize the association between different omics data, enhance the representation of the data and learn the differences and associations between multi-omics data more effectively.

The input dimension of the multi-granularity scanning module is 52, and the sliding window dimension is set to 3. By sampling the features (the sampling step is set to 1), this module generates 18 scanning subsamples. Then, two random forests are used to train the scan subsamples. Since each scan subsample can generate a 2-dimensional probability feature vector after going through random forest A and random forest B, a 72-dimensional probability feature vector is obtained. Finally, the 72-dimensional representation vector corresponding to each gene is spliced and used as the input of the cascaded random forest.

### Deep Forest model scoring

To improve the effectiveness of the features, we use a cascade forest to process the vectors derived from the multi-granularity scans and obtain the results. The cascade forest consists of many layers of forests, each of which includes different kinds of random forests (ordinary random forest and completely random forest), forming a deep learning model similar to a multilayer perceptron. The overall multilayer structure of the cascade forest is shown in Fig.3. In the cascade forest, the first layer of the forest takes the output of the multi-granularity scan as input and can be obtained as follows:

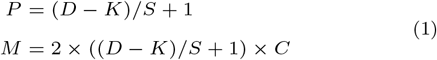

**Fig. 3.**
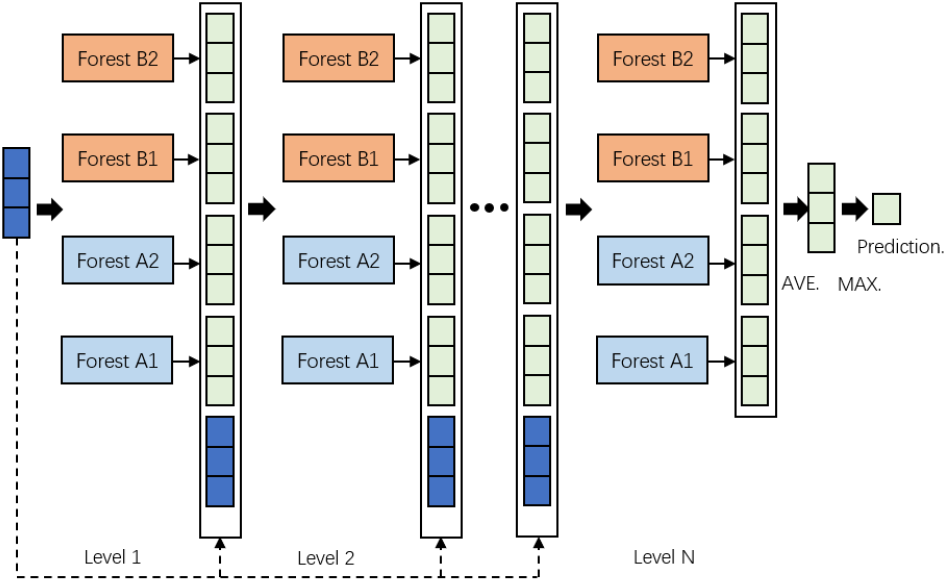
Cascading Forest Module. It uses sliding windows to extract feature vectors, which can enhance the nonlinear characterization ability of the model and maximize the relationship between the order of sample features and the prediction accuracy.

Where D denotes the feature dimension of the original data, K denotes the size of the sliding window, S is the step length of each slide, P is the number of subsamples of K-dimensional features, C denotes the length of the class vector, and M is the output of the multi-granularity scan.

The input of each subsequent layer consists of a combination of the feature vectors obtained from the previous layer and the original input (i.e., the input of the first layer). Each layer in the cascade forest produces a class vector as its output. The probability that each subtree in the cascade forest predicts that sample x belongs to class c can be expressed as:

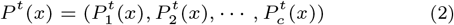

where *t* ∈ *T* and *T* is the number of decision trees in each level of the cascade forest. Each forest obtains a representation of the class C vector 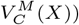 of gene x:

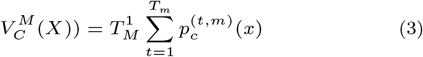

where *m* ∈ *M* and *M* is the number of forests in each layer of the cascade forest. The output of all forests in that cascade forest is obtained as follows:

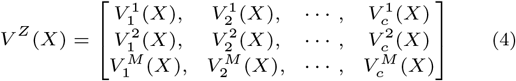

where *z* ∈ *Z* and *Z* is the number of levels in the cascade forest. We then calculated the prediction score of sample x in this cascade forest:

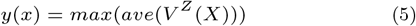

where max() is the maximum function, and ave() is the average function.

The resulting score vector stitched with the result of the multigranularity scan output, which is the original feature vector, as input to the next layer, denoted as:

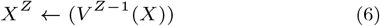

At each extension of a new level, the test set is used to verify the performance of the entire cascade forest, and if there is no significant increase in accuracy, the training is terminated. The final class vector is obtained, the average of the various possibilities is calculated, and the class with the highest value is selected as the final classification result. Due to this property, the number of classes in the cascade is adaptive, unlike the complex and fixed model structure of many deep neural networks. Thus, stable recognition results are obtained even with limited training samples.

### Feature evaluation methods

We interpreted the feature input and the model output with the random forest approach to analyze which omics features are most relevant to cancer druggable genes. We analyzed the contribution of features with the calculation of the Gini scores:

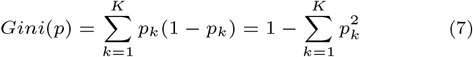

Where k represents the kth category. The score of the node m is calculated as the amount of change in the Gini score before and after branching of m:

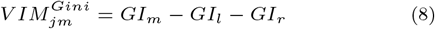

Finally, for the n-class decision trees, the importance scores obtained from all decision trees are normalized as:

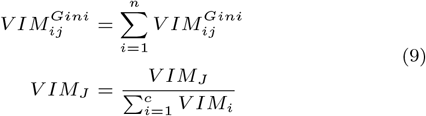

### The cross-validation process

We evaluated the performance of DF-CAGE using the OncoKB dataset, which was downloaded from the Oncology Knowledge Base, a comprehensive precision medical oncology database [30]. The database provides detailed, evidence-based information on somatic and structural mutations to aid treatment decisions. There are 4932 mutations involving 642 genes, 45 tumors, and 89 drugs. The experimental evidence is divided into different levels according to its source and credibility (Level 1 - FDA approved, Level 2A and 2B - the standard of care, Level 3A and 3B - clinical evidence, Level 4 - biological evidence, Levels of Resistance (Level R1-3/ Level R1-2) - drug resistance). This study obtained a total of 45 cancer druggable genes as positive samples for model training by screening genes at Level 3 and above. To verify the stability of DF-CAGE performance under different conditions, we randomly selected a certain number of genes as negative samples in the ratio of 1:1, 1:2, 1:10, 1:20 among 20,000 genes (positive sample data have been removed). The genes in the OncoKB database have very reliable evidence. Using these highly reliable genes for training can guarantee the model’s accuracy to the maximum extent. We used a five-fold cross-validation method to train and test DF-CAGE to prevent overfitting. We divided all the data into five equal parts and then took one of them as the test set each time without repeating and used the other four parts as the training set. To facilitate the performance comparison when the data are unbalanced, we used the AUROC evaluation index to evaluate the performance of each method. Specifically, we calculated the model’s performance on the test data, plotted the ROC curves for five training scenarios, and finally reported the average accuracy of the five-fold verification.

### Validation of DF-CAGE on other datasets

To further validate the power of the DF-CAGE method, we used the TARGET dataset to validate the accuracy of the model. The TARGET (tumor alterations relevant for genomics-driven therapy) [31] dataset was downloaded from the CGA platform, which includes a total of 135 cancer druggable genes that have been used for cancer druggable genes analysis by the previous study. We collected all the 135 cancer druggable genes as the positive samples for the test set and randomly selected among ~20,000 genes in a 1:1 ratio of positive to negative samples as negative samples and combined them into a testing set. We used the model previously trained on the OncoKB dataset (the model with the best performance of the AUROC metric was selected) to predict the TARGET dataset and obtain the probability scoring. Since the ratio of positive and negative samples of the constructed TARGET cancer druggable genes test set is fixed, we used both AUROC and AUPRC performance metrics to measure the performance of DF-CAGE and other methods.

We also used DrugBank, which is primarily used in drug targeting studies, to evaluate the performance of each method. The dataset is downloaded from the DrugBank portal [32] (https://go.drugbank.com/releases/latest). DrugBank is one of the most comprehensive drug databases, but it does not focus on cancer. We screened all drugs with target genes and manually screened 98 cancer druggable genes against the drug descriptions (see Supplementary Table 2 for details).

### The benchmark dataset and comparison methods

We constructd DF-CAGE training and prediction sets according to different positive and negative sample ratios. To evaluate the performance of DF-CAGE and compare the performance on each dataset when the positive and negative samples vary, we used the AUROC value as the evaluation metric on the OncoKB test set. We also used the AUROC value as an evaluation metric on the OncoKB test set (refer to Supplementary Material 3 for calculating the AUROC metric). Due to the high correlation between driver genes and cancer druggable genes, we selected five state-of-the-art driver gene prediction methods as DF-CAGE comparison algorithms: OncodriveCLUST (version 0.4.1), OncodriveFML (version 1.0.2-alpha), MutSig2CV (version 1.0), MuSiC (version 0.2), and e-Driver (version 1.0). Among these methods, OncodriveCLUST, MutSig2CV, and MuSiC rely mainly on somatic mutation frequency-related information for prediction; OncodriveFM and e-Driver are gene function-based algorithms.

## Results

### Validation of DF-CAGE to identify cancer druggable genes on the OncoKB dataset

To illustrate the high reliability, accuracy, and rationality of DF-CAGE in predicting cancer druggable genes, we used the OncoKB dataset as the training dataset for the model. It contains the mutations corresponding to targeted drugs, the biological and oncological impact of mutations, and the frequency distribution characteristics of mutations in the population. Although the sources of information in the knowledge base are diverse, each piece of information is regularly reviewed and revised by the Clinical Genomics Annotation Committee to ensure the accuracy and rigor of the information. Compared with other databases that include somatic mutations, the main content of OncoKB is related to the precise use of drugs in oncology. The knowledge base can be used as the gold standard for identifying drug-available cancer genes.

We collected the ‘allCuratedGenes’ profile from the OncoKB database and extracted all the genes of sensitivity level in the file to obtain 45 cancer druggable genes. These 45 cancer druggable genes were used as a positive sample set. About 70% of the cancer druggable genes in OncoKB were previously reported as cancer drivers. To construct a reasonable negative sample set, we collected the driver genes that could be identified so far, excluding 299 known cancer driver genes among 20,000 human genes and 45 cancer druggable genes in the positive sample set, and randomly selected a different proportion of the remaining of the genes randomly selected in different proportions as well as those found in previous studies (TTN, CACNA1E, COL11A1, DST and other genes unlikely to promote cancer development) as negative samples, and to ensure the reliability of our approach and performance in different situations, the ratios of positive and negative samples were 1:1, 1:2, 1:10, and 1:20, respectively.

Due to the influence of the rare known cancer druggable genes and the unbalanced positive and negative samples, we used the strategy of five-fold cross-validation, the roc evaluation metrics, and the Mann-Whitney test to evaluate the performance of DF-CAGE to prevent the overfitting phenomenon. Because of the lack of specific methods developed for druggable cancer gene prediction and the high correlation between the study of drug-available cancer genes and the study of cancer driver genes, we used the state-of-the-art cancer driver discovery tools as the comparison approaches (see Supplementary Material 4 for details). The performance of each method on the OncoKB dataset is shown in Fig.4.

**Fig. 4.**
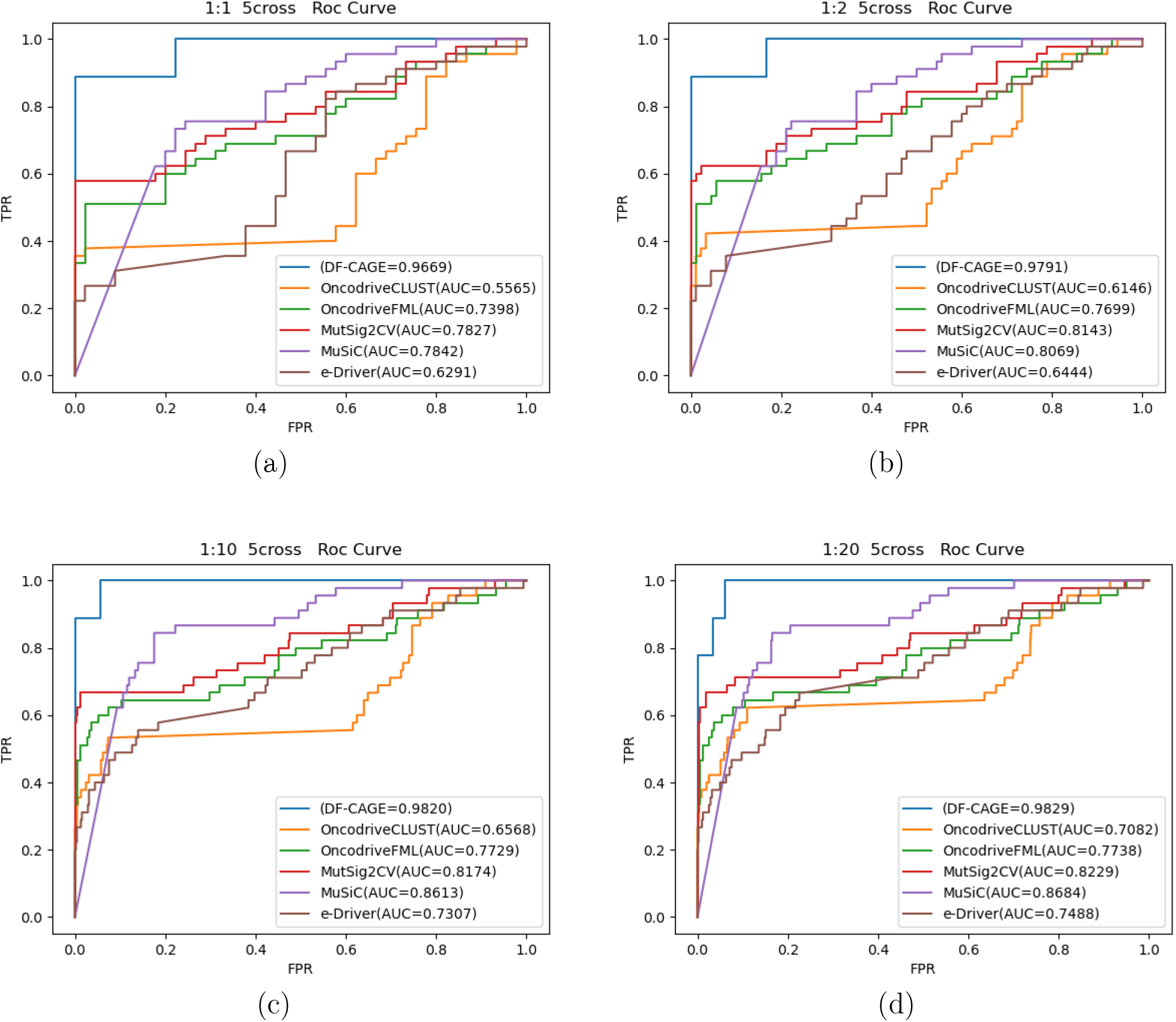
Results of the five-fold cross-validation across four testing scenarios.The ratio of positive and negative samples is (a) 1:1, (b) 1:2, (c) 1:10, (d) 1:20. Performances of DF-CAGE and other approaches including OncodriveCLUST, OncodriveFML, MutSig2CV, MuSic and e-Driver.According to the roc curve, DF-CAGE’s performance is significantly better than the other approaches.

We found that DF-CAGE has the best AUROC performances with a positive to negative sample ratio of 1:1, reaching 0.9669, while we also calculated the Mann-Whitney significant difference check to obtain p-values (p-value=2.92e-14). The next is MuSiC, with an AUROC performance of 0.7842 with a p-value of 1.61e-6. The AUROC performance of OncodriveCLUST, OncodriveFML, MutSig2CV, and e-Driver were 0.5565, 0.7398, 0.7827, and 0.6291, respectively, with p-values of 0.352, 8.99e-5, 3.85e-6, 0.035. We found that DF-CAGE has a notable performance improvement in the AUROC metrics compared with other approaches. Meanwhile, DF-CAGE achieved the most significant p-value in the Mann-Whitney test and the most significant difference between druggable and noncancer druggable genes, indicating that the DF-CAGE method can accurately identify cancer druggable genes. Since cancer druggable genes are relatively rare among all genes, the positive and negative sample ratio setting at 1:1 may not be consistent with the true proportion of cancer druggable genes. For this reason, we continued to construct test datasets with positive and negative sample ratios of 1:2, 1:10, and 1:20. the performance of DF-CAGE improved with the negative samples increased, which may be due to the difference between most of the genes in the negative sample set and the cancer druggable genes significantly, and the difficulty of the task slightly decreased as the number of samples increased. Also, to more visually demonstrate the stability of DF-CAGE performance at different scales, we used a box plot for all the results, as detailed in Supplementary Material 5.

### Verification of the stability of DF-CAGE on the TARGET dataset

To further remove the effect of overfitting on model training and to validate the accuracy and reliability of DF-CAGE, we used the TARGET database under the CGA platform. This dataset contains 135 cancer-related genes with clinical evidence. To prevent overfitting, we held out 45 genes in this test set used for DF-CAGE model training but appeared in the test dataset.

We extracted all the genes from the TARGET database and used them as the positive sample set for the test set. Similarly, the negative sample set was randomly selected from the other genes (45 genes used for previous model training were held out). We take the same number of negative samples as the positive samples to construct the test dataset and compare the performance. We used DF-CAGE to score all genes in the TARGET test set probabilistically using the model with the best performance of the AUROC metric on the OncoKB dataset and compared the performance of the DF-CAGE model with the comparison algorithms. To reflect the performance of our model more comprehensively, we used both pr and roc metrics and the experimental strategy of Mann-Whitney significant difference check on this balanced dataset and performed enrichment analysis of various methods on the TARGET dataset (Fig.5).

**Fig. 5.**
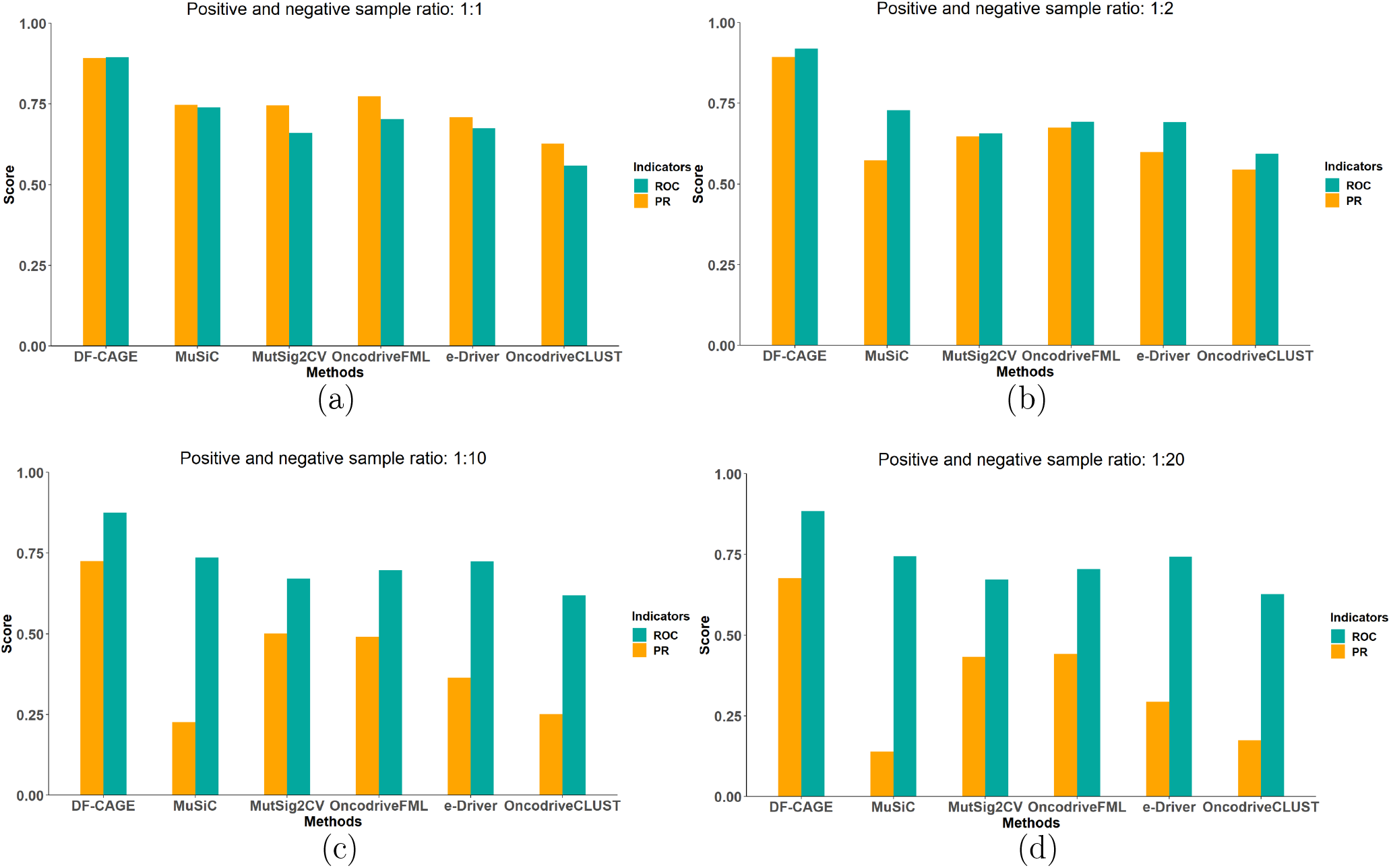
The performance of different methods across four test scenarios of the TARGET dataset.The ratio of positive and negative samples is (a) 1:1, (b) 1:2, (c) 1:10, (d) 1:20.

It is intuitive to see that the DF-CAGE model significantly outperforms the other methods in different ratios. While the ratio of positive and negative samples is 1:1, the AUPRC of DF-CAGE method is 0.8917, and the AUROC is 0.8933, while the Mann-Whitney test p-value is 5.00e-20, outperformed OncodriveCLUST (AUPRC=0.6270, AUROC=0.5585, p-value=1.67e-1), Oncodriver-FML (AUPRC= 0.7736, AUROC= 0.7020, p-value=2.48e-06), MutSig2CV (AUPRC= 0.7454, AUROC= 0.6595, p-value=2.02e-4), MuSiC (AUPRC= 0.7468, AUROC= 0.7392, p-value=2.05e-08), e-Driver (AUPRC= 0.7082, AUROC= 0.6743, p-value= 4.86e-05). The results for other positive and negative sample proportions were similar to the above (see Supplementary Material 5 for specific analysis). The model parameters and the classification thresholds were based on the training process. We also performed the enrichment analysis on the TARGET dataset (excluding cancer druggable genes in the oncoKB dataset, leaving 91 cancer druggable genes). We used Fisher’s exact test to calculate empirical P-values. While the ratio of positive and negative samples is 1:1, the DF-CAGE method reported 29 cancer druggable genes according to the classification threshold (Fisher’s exact test p-value =1.55e-8). the number of the TARGET genes reported by other methods was: OncodriveCLUST (1), OncodriveFML (20), MutSig2CV (16), MuSiC (28), e-Driver (6).The performance of enrichment analysis of other methods is not as well as that of DF-CAGE: OncodriveCLUST (Fisher’s exact test p-value = 4.99e-1), OncodriveFML (Fisher’s exact test p-value =2.95e-7), MutSig2CV (Fisher’s exact test p-value =7.41e-6), MuSiC (Fisher’s exact test p-value 2.80e-08), and e-Driver (Fisher’s exact test p-value =1.43e-2) (see Supplementary Material 6 for other test sets). Although all genes in the TARGET test set did not appear in the DF-CAGE training data, DF-CAGE can also pinpoint cancer druggable genes accurately in the TARGET dataset, indicating that the DF-CAGE model can reliably and accurately identify cancer druggable genes.

In addition, to further validate the reliability and accuracy of the DF-CAGE model, we also conducted evaluations on the DrugBank cancer druggable genes set. Compared to other comparative methods, DF-CAGE also achieved excellent performance (See Supplementary Material 7 for details).

### DF-CAGE report 465 potentially cancer druggable genes

The evaluation of DF-CAGE on multiple datasets described above has demonstrated that DF-CAGE can accurately predict cancer druggable genes. Therefore, we expected to use DF-CAGE to explore the entire genome of 20,000 coding region genes to find novel cancer druggable genes, which will help us understand the relationship between cancer druggable genes and multi-omics molecular data (such as the mutation data). We used 135 cancer druggable genes from the TARGET dataset as the DF-CAGE training set, and to prevent overfitting and to evaluate model performance, we trained the models using a five-fold cross-validation approach and used the model with the best performance on the training set to make predictions on all 20,000 human coding genes. After we obtained the DF-CAGE scores for all coding genes, we used the best score threshold obtained during cross-validation (cutoff = 0.55) to obtain 465 potential genes that constitute the cancer druggable genes (CDG-genes) set.

### Explore the relationship between the cancer druggable genes and the CGC genes

Previous studies have mainly analysed the pathogenic mechanisms of driver genes to identify cancer druggable genes among them. In this study, we directly used DF-CAGE to obtain a set of 465 druggable cancer genes (CDG-genes). We analysed this gene set and the cancer driver gene set (Fig.6) and found that a total of 215 genes out of the 465 genes predicted by DF-CAGE also appeared in the CGC database, which suggested that there are still a large number of driver genes that are still druggable, only that the complementary therapies are yet to be discovered. For instance, KRAS is a well-known driver gene in colorectal cancer, which was considered a highly reliable druggable gene by DF-CAGE and was once considered non-druggable [33, 34]. However, Gongmin Zhu et al. developed a KRAS G12C allele-specific inhibitor, which breaks the traditional view that KRAS is not druggable, changes the therapeutic outlook of KRAS-driven tumors, and offers the possibility to overcome KRAS mutations in CRC.

**Fig. 6.**
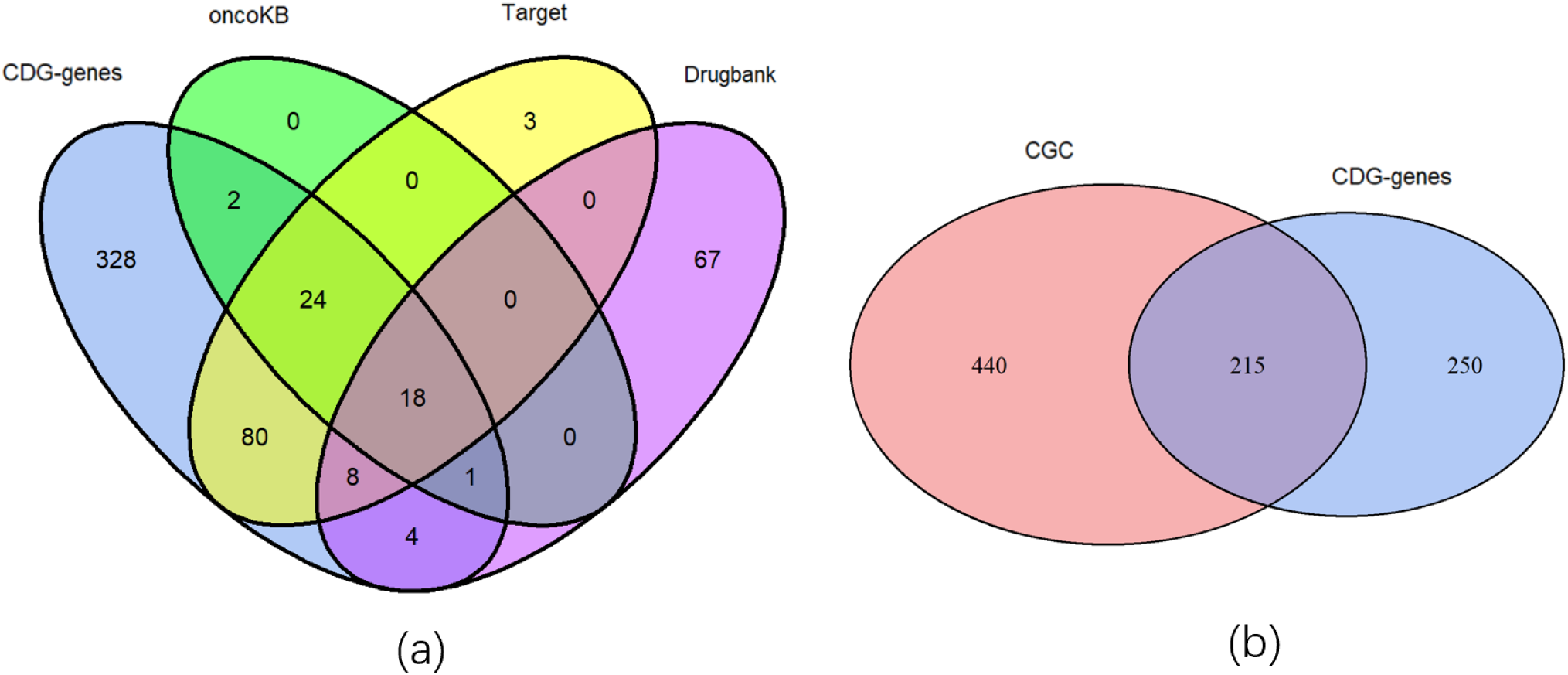
(a) Intersection of CDG-genes and OncoKB, TARGET, DrugBank druggable gene sets (plot using the R language toolkit ‘ggvenn’). (b) Intersection of CDG-genes and CGC driver genes.

We also found that DF-CAGE predicted 250 genes outside of CGC, which contained highly reliable, reliable, and potential genes. For instance, DF-CAGE reported ROS1, Tex10 that are not in the CGC gene set. The ROS1 gene has been reported as a drug-targeted to treat lung cancer [35, 36, 37] and is a highly reliable gene that has been used in clinical medicine. In addition, studies have shown that Tex10 plays a decisive oncogenic role in HCC tumorigenesis by activating the STAT3 signaling pathway and promoting chemoresistance to maintain cancer stem cell properties. Therefore, targeting Tex10 may provide a novel and effective therapeutic strategy to inhibit the tumorigenicity of advanced HCC [38, 39, 40]. The above results suggest that DF-CAGE helps find cancer druggable genes beyond the known driver genes, which have received relatively little attention in previous studies.

### Analysis of the effectiveness of omics data for identifying cancer druggable genes

The above analysis suggested that our method DF-CAGE has high accuracy, reliability, and stability in identifying cancer druggable genes, exceeding other driver genes identification methods such as Mutsig2CV and ActiveDriver. The features used in DF-CAGE include genomics data (somatic mutations, CNVs), epigenetic data (genome-wide DNA methylation), and transcriptomic data (mRNA-seq), while other methods mainly use mutation-related features. Multi-omics analysis can more comprehensively characterize all aspects of genes, learn the common patterns among different omics data, and reduce the noise.

On the task of direct druggable gene identification using multiomics data, none of the previous studies have been done to assess the contribution of each omics data to the identification results. To understand the association between CDG-genes and input features and obtain which omics data are most relevant to cancer druggable genes, we performed the model interpretation analysis on the integrated multi-omics data with random forest. We ranked the Gini scores for all the input features (Fig. 7).

**Fig. 7.**
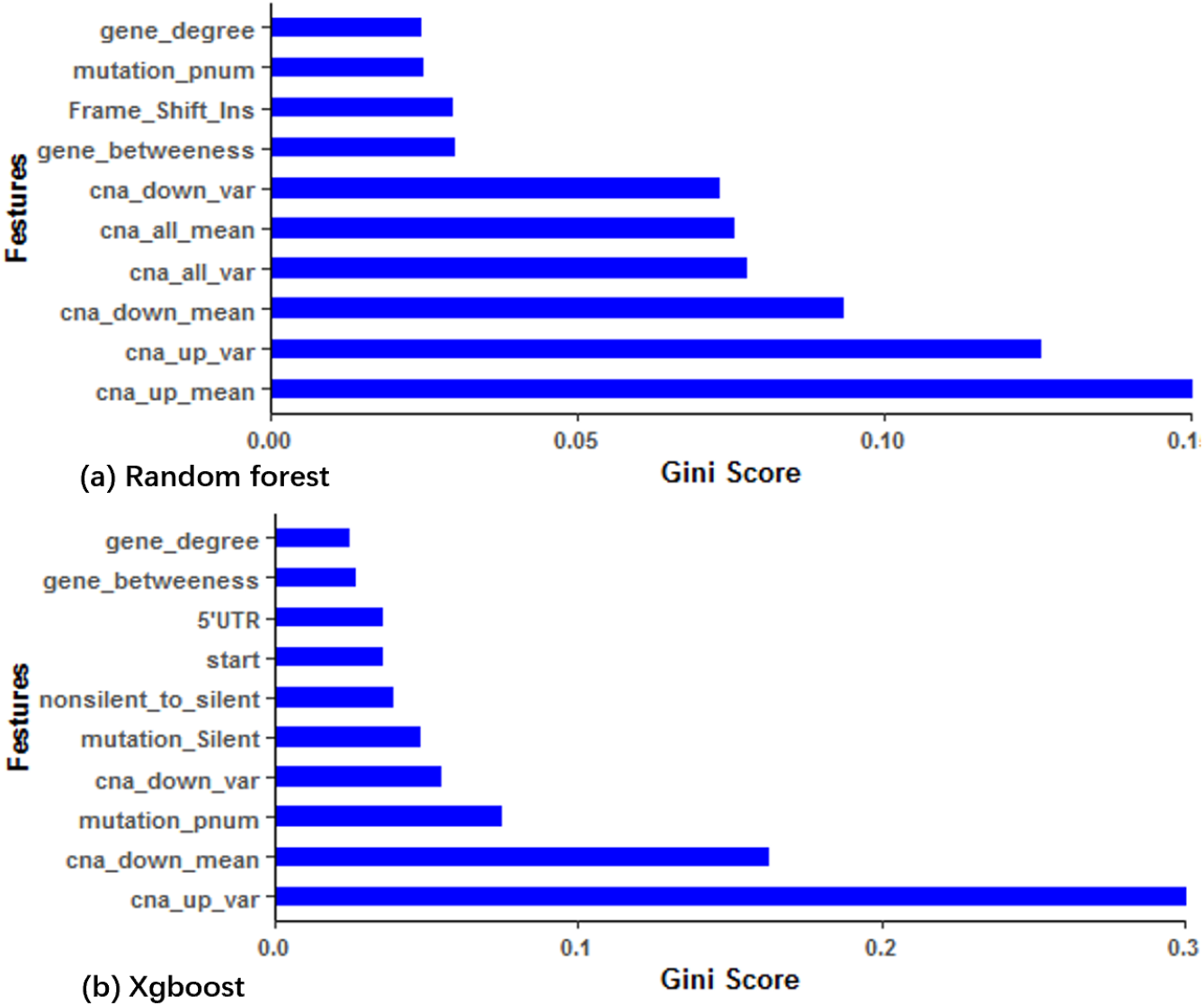
The feature importance analysis results of DF-CAGE: (a) using random forest. (b) using Xgboost.

Among the ten most critical multi-omics-related features of DF-CAGE, the top6 features are associated with CNV data: ‘up_mean’ (mean of copy number increase, Gini score =0.0934), ‘all_mean’ (mean of CNAs, Gini score= 0.0803, ‘down_mean’ (mean of copy number decrease, Gini score =0.0777, ‘all_var’ (variance of CNAs, Gini score=0.0688), ‘down_var’ (variance of copy number decrease, Gini score=0.0639, and ‘up_var’ (variance of copy number increase, Gini score=0.0584). Unlike the identification of cancer driver genes dominated by somatic mutation-related features, cancer druggable genes are most strongly associated with CNV-related features. We also found that the stronger the interconnectivity between genes, the more likely they are druggable. Among the top10 features, the Gini score of ‘gene_degree’ (number of linked genes) was 0.0529, and the Gini score of ‘gene_betweeness’ (intergenic centrality) (14) was 0.0486. The results indicate that genes strongly associated with other genes are more likely to be targets of cancer therapeutic drugs. The feature of ‘pnum’ (number of mutations, Gini score=0.0688) ranked 10th in the 52-dimensional omics input, which is also essential for predicting cancer druggable genes, indicating that cancer druggable genes tend to be enriched in genes with high mutation rates in the population. To ensure that the feature importance did not change substantially due to the change in method, we also used another model interpretation method: xgboost [41] and analysed the importance of the above features (Fig.7(b) and Supplementary Material 8). We found that the top-ranked features of the two methods were very similar, which demonstrates that the order of importance of features is stable.

## Discussion

This paper proposes DF-CAGE, a machine learning-based cancer druggable gene discovery method. DF-CAGE integrates multiple omics data (DNA methylation, mRNA-seq, CNVs, and somatic mutations) and performs druggable gene discovery. Through the multi-granularity scanning module, DF-CAGE learns the association between different omics data and enhances the representation of the data to understand the differences and associations between multiple omics data. DF-CAGE characterizes the non-linear feature inputs through the cascade forest module to get the output druggable gene classification results. To the best of our knowledge, DF-CAGE is the first machine learning method to identify cancer druggable genes based on multi-omics data integration. The performance of DF-CAGE exceeds AUROC of 0.9 on both OncoKB and TARGET datasets, and the crossvalidated AUROC even reaches 98% on the OncoKB dataset. Unlike the traditional cancer driver gene discovery approaches (such as MutSig2CV, ActiveDriver, OncodriveFM, MuSiC), DF-CAGE directly reported the candidate druggable genes. We obtained a druggable gene dataset, CDG-genes, with DF-CAGE, including 465 genes. We validated a significant number of genes in the CDG-genes set with the literature investigation and classified these genes into 130 highly reliable genes, 31 reliable genes, and 304 potential genes(See Supplementary Material 9 for a more detailed discussion).

The cognition of whether a gene is cancer-druggable is inseparable from biomedical technology development. With the development of pharmaceutical technology in the future, DF-CAGE will identify more cancer druggable genes while new genes are introduced into its model training set. The relationship between driver genes and cancer druggable genes will become more prominent. Our future work includes exploring and identifying cancer druggable genes with the corresponding drugs on most individual cancer types (especially on some rare cancer types). It is a more challenging task since the ground truth is limited. The latest machine learning techniques, such as the few-shot learning and meta-learning methods, will be considered to accomplish cancer type-specific druggable gene discovery.

## Data Availability

The TCGA pan-cancer dataset,the OncoKB dataset,the TARGET dataset and the DrugBank portal used by DF-CAGE are available on http://gdc.caner.gov/aboutdata/publications/pancanatlas, https://www.oncokb.org/, https://software.broadinstitute.org/cancer/cga/target and https://go.drugbank.com/releases/latest. The source codes, the scoring results of DF-CAGE across the benchmark data sets and CDG-genes dataset (including three tables of highly reliable genes, reliable genes and potential genes) are available at https://github.com/haiyangLab/DF-CAGE.

## Competing Interests

All of the authors declare no competing financial interests.

## Author Contributions Statement

H.Y. and Z.W. designed the study. L.G. developed the DF-CAGE framework and implemented it. C.R., D.L., J.Z., Z.W. collated multi-omics data and completed the performance comparison. H.Y. and L.G. analyzed the experimental results. H.Y. and L.G. wrote the manuscript. All authors read and approved the manuscript.

### Key points

- This study proposes DF-CAGE, a novel machine learning method for cancer-druggable gene discovery. DF-CAGE model consists of two main components: the multi-granularity scan module and the cascade forest module. With the multi-granularity scanning module, DF-CAGE can learn the association between different omics data, enhancing the representation of the data and learning the differences and associations between multi-omics data more effectively. It uses sliding windows to extract feature vectors, which can enhance the nonlinear characterization ability of the model and maximize the relationship between the order of sample features and the prediction accuracy.
- DF-CAGE integrated the somatic mutations, copy number variants, DNA methylation, and RNA-Seq data across 10000 TCGA profiles to identify the landscape of the cancer-druggable genes. we analyzed the contribution of the omics data to the identification of druggable genes. We found that DF-CAGE reports druggable genes mainly based on the CNAs, the gene rearrangements, and the mutation rates in the population. Multi-omics cancer data has the potential to discover new cancer therapeutic targets. These findings may enlighten the study and development of new cancer drugs.
- We found that DF-CAGE discovers the commonalities of currently known cancer-druggable genes from the perspective of multi-omics data and achieved excellent performance on OncoKB, Target, and Drugbank data sets (AUROC>0.8).
- DF-CAGE obtained a druggable gene dataset, CDG-genes, including 465 genes. We validated a significant number of genes in the CDG-genes set with the literature investigation and classified these genes into 130 highly reliable genes, 31 reliable genes, and 304 potential genes.

## Acknowledgments

We are grateful to Dr. Tianhao Gu, Dr. Wei Guo, Dr. Mengping Yang for the critical reading of the manuscript and the other members of the AIMC lab for practical discussions and suggestions.

## Funding

This work is supported by Natural Science Foundation of China under Grant No. 61902126, Shanghai Science and Technology Program ‘‘Distributed and generative few-shot algorithm and theory research” under Grant No. 20511100600, Shanghai Science and Technology Program ‘Federated based cross-domain and cross-task incremental learning” under Grant No. 21511100800.

**Hai Yang** is an associate professor in the Department of Computer Science and Engineering at East China University of Science and Technology. His research topics include artificial intelligence, machine learning, big data, and bioinformatics.

**Lipeng Gan** is a master student in the Department of Computer Science and Engineering at East China University of Science and Technology. His research interests include machine learning and bioinformatics.

**Rui Chen** is a research lecturer in the Department of Molecular Physiology and Biophysics at Vanderbilt University, where his research topics include molecular physiology and biophysics.

**Dongdong Li.** is an associate professor at the Department of Computer Science and Engineering at East China University of Science and Technology. Her research topics include machine learning, artificial intelligence, affective computing, and multimedia computing.

**Jing Zhang** is an associate professor in the Department of Computer Science and Engineering, East China University of Science and Technology. Her research topics include image semantic content analysis, video content structure analysis, ontology-based video image annotation and retrieval.

**Zhe Wang** is a professor at the Department of Computer Science and Engineering at East China University of Science and Technology. His research topics include pattern recognition, machine learning and its applications, data mining and big data analysis, image processing and analysis, bioinformatics.

## Notes

### Competing Interest Statement

The authors have declared no competing interest.

