## Supplementary material for "From multi-omics data to the cancer druggable gene discovery: a novel machine learning-based approach"

**Supplementary materials**

**Supplementary Note 1: Data pre-processing and feature selection**

First, to process the somatic mutation data, we obtained the somatic mutations data from PanCancerAtlas and used the column of 'Variant_Classification' (3'Flank, 3'UTR, 5'Flank, 5'UTR, 'Frame_Shift_Del', 'Frame_Shift_Ins', 'In_ Frame_Del', 'In_Frame_Ins', Intron,' Missense_Mutation', 'Nonsense_Mutation', 'Nonstop_Mutation', ‘RNA’, ‘Silent’, 'Splice_Site', 'Translation_Start_Site') and the column of Variant_Type (DEL, INS, ONP, SNP, TNP) to construct the mutation-related features. We counted the number of various types of mutations corresponding to each gene as the feature values. While a gene does not contain a certain type of mutation, its corresponding feature value is set to 0.

Next, we downloaded the RNA-seq data from PanCancerAtlas. We focused on the ~20,000 protein-coding genes and calculated the mean and variance of the gene expressions across the TCGA samples as the corresponding features. we used the other genes' mean value on RNA-seq

to fill in the missing data.

For the CNV data, we collected the relevant profiles from the TCGA dataset. After that, we calculated the mean and variance of the CNVs occurring on each gene. To consider both aberrations deletions of CNVs, we counted the mean and variance in these two cases separately. Like the case encountered for mRNA data, we fill in the missing data in the feature matrix by calculating the mean values.

We obtained DNA Methylation (450K Only) data from PanCancerAtlas. Similar to the RNA data, the mean and variance of methylation for each gene were calculated, except that the data were not log-transformed, and the DNA methylation data were mapped to specific genes by using the mapping table of the ‘CG number’ and the ‘gene id.’

We introduced some other features, including the gene degree of the BioGrid's network (number of linked genes), intergenic centrality, and genomic covariate data (including expression level, DNA replication time, and Hic (chromatin state of genes). Finally, we obtained 52-dimensional features as model inputs for DF-CAGE based on multi-omics data.

**Supplementary Note 2: The model training process of DF-CAGE**

DF-CAGE is a deep learning approach that uses an ensemble learning method based on multi-layer forests. It mainly consists of the multi-granularity scanning module and the cascade forest module. DF-CAGE especially has the following characteristics: in the deep forests, it requires fewer hyperparameters to be fitted and is not prone to overfitting even if the model is trained with a small amount of data; the complexity of the model can be determined automatically by correlating it with the data, and the model can be implemented without using backpropagation.

An intuitive way to improve the performance of the deep forest is to increase its model complexity. Some important parameters determine the model complexity of deep forest in DF-CAGE: ‘n_estimators’ (specify the number of estimators in each cascaded layer), ‘n_trees’ (specify the number of trees in each estimator). Using the above larger parameters, the performance of deep forest may improve the performance of complex datasets, while the training set size of the DF-CAGE method is small. Therefore, we set the above three parameters as follows: ‘n_estimator’ = 2, ‘n_trees’ = 100, ‘max_layers’=20. To get better accuracy of DF-CAGE, the parameters ‘use_predictor’ (decide whether to use the predictor connected to the deep forest), ‘predictor’ (specify the type of predictor, should be one of ‘forest,’ ‘xgboost,’ ‘ lightgbm’) are set to ‘False’ and ‘forest,’ respectively.

The training process of DF-CAGE is through 52-dimensional multi-omics features. Firstly, the multi-granularity scanning module is adopted to enhance the cascade forest. Then, sampling is performed using sliding windows of multiple sizes to obtain more feature subsamples. This paper’s default sliding window sizes are 3, 6, 12. When the sliding window size is 3, 49 3-dimensional sub-features can be obtained by inputting 51-dimensional features. The random forest can obtain the 49 2-dimensional class vectors. Finally, the 196 feature vectors are obtained by stitching, and the other sliding windows are similar. As shown in Fig.S1, the feature vectors obtained in the multi-granularity scanning stage were input into the cascade forest module. Each cascade layer had four random forests, and each layer got 8-dimensional feature vectors, which were used as the input of the next cascade layer to enhance the original features. To reduce the risk of overfitting, the class vectors of each forest are generated through k-fold cross-validation. Each sample is trained k-1 times as training data, resulting in k-1 class 2-dimensional vectors, which are then averaged to be the final feature vectors of this forest. Then, the 2D feature vectors of the four forests are linked together as the enhanced feature vectors of the next layer. Finally, we obtained the identification results of DF-CAGE.

**Supplementary Note 3: The evaluation metrics on the test sets**

Firstly, the confusion matrix elements in the following form were defined: TP was defined as the number of actual cancer druggable genes identified to be cancer druggable genes; FP was defined as the number of actual non-cancer druggable genes identified to be cancer druggable genes; FN was defined as the number of actual cancer druggable genes identified to be non-cancer druggable genes; TN was defined as the number of actual non-cancer druggable genes identified to be non-cancer druggable genes. According to the above confusion matrix elements, accuracy, precision, F1 value, and AUC (Area Under Curve) were used to evaluate the performance of identified cancer druggable genes. The accuracy rate represents the probability that the entire sample, including cancer druggable genes and non-cancer druggable genes, is correctly identified and is calculated as:

$$\begin{aligned} A_{c}=\frac{TP+TN}{TP+TN+FP+FN}\#\left( 1 \right) \end{aligned}$$

The precision rate represents the probability that the identified cancer druggable genes are cancer druggable genes, and the calculation method is as follows:

$$\begin{aligned} P_{r}=\frac{TP}{TP+FP}\#\left( 2 \right) \end{aligned}$$

The F1 value index can comprehensively evaluate the precision rate and recall rate of cancer druggable genes identification methods, and the calculation method is as follows:

$$\begin{aligned} F_{1}=\frac{P_{r}\times R_{e}}{P_{r}+R_{e}}\#\left( 3 \right) \end{aligned}$$

Where R represents the recall rate of the identification model, which is defined as:

$$\begin{aligned} R_{e}=\frac{TP}{TP+FN}\#\left( 4 \right) \end{aligned}$$

The AUC represents the area under the receiver operating characteristic (ROC) curve, and the value is between 0 and 1. The larger the index, the better the identification performance of the model.

**Supplementary Note 4: Exploring the stability of the DF-CAGE model on the OncoKB dataset**

To show the stability of DF-CAGE performance more intuitively under different scales, we investigated the performance of all methods on the OncoKB dataset (The ratio of positive and negative samples is: 1:1, 1:2, 1:10, 1:20). The construction of positive and negative training samples is shown in the main text. Due to the correlation between driver genes and drug-available genes, we selected five state-of-the-art driver gene identification methods as the DF-CAGE comparison algorithms: OncodriveCLUST, OncodriveFML, MutSig2CV, MuSiC. We used the 5-fold cross-validation to obtain the DF-CAGE scores on the OncoKB dataset. Other methods used the scores from the previous TCGA study. In addition, we used the AUROC metric to evaluate performance:

From Table S1, we can intuitively see that the average performance of DF-CAGE is higher than other methods when the positive and negative samples are 1:1, 1:2, 1:10, and 1:20. The upper and lower AUROCs of DF-CAGE are 0.99 and 0.96. The upper and lower AUROCs of MuSiC are 0.87 and 0.77. The upper and lower AUROCs of OncodriverFML are 0.78 and 0.72. Musig2cv has upper and lower AUROCs of 0.83 and 0.77, and OncodriveCLUST has upper and lower AUROCs of 0.71 and 0.53. Thus, it is shown that in different proportions of positive and negative samples case, DF-CAGE outperforms other methods in terms of performance and stability.

**Supplementary Note 5: Analysis of the results of other machine learning algorithms on the OncoKB dataset**

To fully illustrate the superiority of the performance of the deep forest method, we also compared the deep neural network (DNN), SVM, and the Random Forest algorithm method with our approaches on the OncoKB dataset (based on the same model input of DF-CAGE). To ensure the correctness of the comparative experiments, we adopt the same benchmark strategy of 5-fold cross-validation and make comparisons according to the ratio of positive and negative samples 1:1, 1:2, 1:10, and 1:20. The benchmark results are shown in the figure below.

Firstly, we set parameters for three machine learning methods (SVM, DNN and Random Forest). The DNN method consists of two fully connected layers with the ReLU function as the activation function, and the Adam as the optimizer. The ‘batch_size’ parameter was set to 32. The SVM method uses the default parameter settings, with the penalty parameter C = 1.0, kernel = "rbf" and gamma ="auto". The Random Forest method uses the default parameter settings and ‘n_estimators’ was set to 30. It can be seen intuitively from Fig.S2 that when the ratio of positive and negative samples is 1:1, the AUROC performance of our method DF-CAGE is 0.9969, and the AUROC performance of other methods are: SVM (0.7477), DNN (0.8059), Random forest (0.9586). The DF-CAGE method has a relatively large improvement compared to SVM and DNN, and the absolute performance of DF-CAGE is improved by ~1% compared to Random Forest. The benchmark results of other positive and negative sample ratios are similar. With the increase of negative samples, the performance of these methods has improved. The AUROC methods of SVM and DNN methods have improved significantly by about 0.88, while for Random Forest, in positive and negative samples, in the case of a ratio of 1:20, the AUROC performance has declined. In summary, the DF-CAGE method can more accurately and reliably predict the available essential genes under different proportions of positive and negative samples.

**Supplementary Note 6: The performance comparison details on the** **TARGET dataset**

For the enrichment analysis of the approaches on the TARGET dataset, we used Fisher’s exact test to calculate the p-values and performed the -log transformation to get the performance of each method (-log10 p-value) (Fig.S3).

While we trained the DF-CAGE model with five-fold cross-validation on the training dataset, the optimal model parameters and the classification thresholds were obtained. We performed enrichment analysis on the TARGET dataset (excluding the cancer druggable genes in the OncoKB dataset, leaving 91 cancer druggable genes). We used Fisher’s exact test and calculated the corresponding p-values. According to the classification threshold, the DF-CAGE method reported a total of 31 TARGET genes with a positive to negative sample ratio of 1:1, and the number of the TARGET genes reported by other methods was: OncodriveCLUST (1), OncodriveFML (20), MutSig2CV (16), MuSiC (28), e-Driver (6). We found that the DF-CAGE was the most enriched on the TARGET data set (P-value=1.55e-08), followed by MuSiC (P-value=2.80e-08), OncodriveFML (P-value=2.95e-07), MutSig2CV (P-value=7.41e-06), e-Driver (P-value=1.43e-2), OncodriveCLUST (P-value=4.99e-1).

Since cancer druggable genes are relatively rare among all genes, the positive and negative sample ratio setting at 1:1 may not be consistent with the true proportion of cancer druggable genes. In the case of positive and negative samples 1:2, the DF-CAGE method reported a total of 54 cancer druggable genes, and the number of the genes reported by other methods was: OncodriveCLUST (1), OncodriveFML (20), MutSig2CV (16), MuSiC (29), e-Driver (6). The DF-CAGE was the most enriched on the TARGET data set (P-value=2.43e-28), followed by OncodriveFML (P-value=6.13e-11), MutSig2CV (P-value=8.96e-09), MuSiC (P-value=1.06e-08), e-Driver (P-value=1.22e-3), OncodriveCLUST (P-value=3.33e-1).

Similarly, in the 1:10 case, a total of 45 cancer druggable genes were reported by the DF-CAGE method, and the number of the genes reported by other methods was: OncodriveCLUST (0), OncodriveFML (20), MutSig2CV (16), MuSiC (29), e-Driver (5). The DF-CAGE was the most enriched on the TARGET data set (Fisher’s exact test P-value =2.33e-47). The performance of enrichment analysis (Fisher’s exact test) of other methods is as follows: OncodriveFML (P-value=1.88e-22), MutSig2CV (P-value=6.04e-18), MuSiC (P-value=2.13e-12), e-Driver (P-value=5.60e-06), OncodriveCLUST (P-value=1.0). In the case of positive and negative samples 1:20, DF-CAGE method reported a total of 45 cancer druggable genes. The number of the genes reported by other methods was: OncodriveCLUST (0), OncodriveFML (20), MutSig2CV(16), MuSiC (32), e-Driver(6). We found that the gene set reported by DF-CAGE was the most enriched (Fisher’s exact test) on the TARGET data set (P-value=1.48e-54), followed by MutSig2CV (1.83e-22), MuSiC (6.33e-14), e-Driver (9.93e-09), OncodriveFML (1.04e-01), OncodriveCLUST (P-value=1.0).

In addition, we used the Mann-Whitney test to show that whether the classification results obtained by each method are significant on the TARGET dataset (Fig.S4). In the case of positive and negative samples 1:1, DF-CAGE achieved the best performance (p-value=5.00e-20), followed by MuSiC (p-value=2.05e-08), OncodriverFML (p-value=2.48e-06), e-Driver (p-value=4.86e-05), MutSig2CV (p-value=2.02e-4), and OncodriveCLUST (p-value=1.67e-1). In the case of positive and negative samples 1:2, DF-CAGE also obtained the best performance (p-value=1.40e-29), MuSiC (p-value=6.80e-10), OncodriverFML (p-value=2.38e-07), e-Driver (p-value=2.43e-07), MutSig2CV (p-value=2.62e-05), OncodriveCLUST (p-value=9.93e-03). In the case of positive and negative samples 1:10, the p-values of all methods are DF-CAGE (1.59e-33), MuSiC (1.02e-13), e-Driver (1.69e-12), OncodriverFML (6.84e-10), MutSig2CV (7.83e-08), OncodriveCLUST (8.12e-05), In the case of positive and negative samples 1:20, the p-values of all methods are DF-CAGE (1.58e-44), MuSiC (p-value=2.75e-15), e-Driver (p-value=4.04e-15), OncodriverFML (3.87e-11), MutSig2CV (2.86e-08), OncodriveCLUST (1.56e-05). We found that the performance advantage of DF-CAGE is considerably higher than other algorithms on four different ratios. This indicates that the DF-CAGE method can accurately and reliably predict druggable genes, and that there is a significant difference between druggable and non-druggable genes identified by the DF-CAGE method.

**Supplementary Note 7: Enrichment analysis of DF-CAGE on the Drugbank dataset**

Simultaneously, on the Drugbank dataset, we performed enrichment analysis for all the methods (DF-CAGE, OncodriveCLUST, OncodriveFML, MutSig2CV, MuSiC, e-Driver). We used Fisher’s exact test, obtained the corresponding P-values, and performed the -log transformation on the results to obtain the performance of each method (-log10 p-values) (Fig.S5).

In the case of 1:1 positive and negative samples, the performance of different methods on the Drugbank dataset are DF-CAGE (2.13e-10), MuSiC (1.32e-04), MutSig2CV (2.67e-03), OncodriveFML (1.54e-01), e-Driver (3.24e-01), OncodriveCLUST (1.65e-01). We can see that DF-CAGE achieved the best performance on the Drugbank data set. Similarly, in the case of 1:2, the performance of all the methods: DF-CAGE (5.01e-16), OncodriveCLUST (3.65e-02), OncodriveFML (6.32e-02), MutSig2CV (4.40e-05), MuSiC (4.16e-05), e-Driver (1.37e-02). In the case of 1:10, the performances of all the methods are DF-CAGE (2,45e-25), OncodriveCLUST (3.99e-04), OncodriveFML (4.52e-05), MutSig2CV (9.52e-10), MuSiC (6.51e-07), e-Driver (3.53e-04). In the case of 1:20, the performances of all the methods are DF-CAGE (4.45e-30), OncodriveCLUST (5.36e-06), OncodriveFML (5.78e-06), MutSig2CV (1.21e-13), MuSiC (7.75e-08), e-Driver (4.64e-05). In the case of different ratios, the DF-CAGE method has the best enrichment performance on the Drugbank dataset. The DF-CAGE has a significant gap compared with other methods, followed by MutSig2CV, and the OncodriveCLUST method has the lowest power. Our method DF-CAGE can reliably and accurately discover cancer druggable genes on the Drugbank dataset.

Similarly, we used the Mann-Whitney test to show that whether the classification results obtained by each method are significant on the TARGET dataset (Fig.S6). In the case of positive and negative samples 1:1, DF-CAGE achieved the best performance (p-value=6.86e-14), followed by OncodriveCLUST (p-value=9.64e-08), MuSiC (p-value=7.00e-07), MutSig2CV (p-value=3.82e-06), e-Driver (p-value=5.08e-05), , and OncodriveCLUST (p-value=1.06e-01). In the case of positive and negative samples 1:2, DF-CAGE also obtained the best performance (p-value=4.22e-18), OncodriveCLUST (p-value=2.10e-09), MutSig2CV (p-value=2.33e-07), MuSiC (p-value=4.56e-07), e-Driver (p-value=1.52e-05), OncodriverFML (p-value=1.85e-02). In the case of positive and negative samples 1:10, the p-values of all methods are DF-CAGE (5.16e-26), OncodriveCLUST (6.83e-14), MutSig2CV (5.73e-10), MuSiC (8.23e-10), e-Driver (5.63e-09), OncodriverFML (2.29e-03). In the case of positive and negative samples 1:20, the p-values of all methods are DF-CAGE (1.84e-28), OncodriveCLUST (p-value=3,94e-16), MuSiC (p-value=4.22e-11), MutSig2CV (p-value=5.57e-11), e-Driver (p-value=1.11e-10), OncodriverFML (p-value=8.93e-04). We found that the performance advantage of DF-CAGE is obvious, which indicated that the DF-CAGE method can accurately and reliably predict druggable genes, and that there is a significant difference between druggable and non-druggable genes identified by the DF-CAGE method.

**Supplementary Note 8: Multi-omics feature importance assessment based on the Xgboost method**

We further analyzed the interpretability of the DF-CAGE model with the evaluation of the feature importance of the multi-omics data on the TARGET dataset. We used the Xgboost algorithm to calculate the contribution ratio of different features to the recognition results. The top10 essential features are shown in Fig.S7 (the importance order of all features is shown in Fig. S8).

We found that CNV-related features are critical in identifying cancer druggable genes. The top10 important features include three CNV-related features. The Gini scores of ‘cna_up_var’ (the variance of all CNVs), ‘cna_down_mean’ (the mean of the duplication of CNVs), and ‘cna_down_var’ (the variance of deletion of CNVs) are 0.3009, 0.1633, 0.0553, respectively. The Gini scores of other top10 features: ‘mutation_pnum’ (number of mutations): 0.0749, ‘mutation_silent’ (silent mutation): 0.0480, ‘nonsilent_to_silent’ (non-silent mutation and silent ratio): 0.0392, ‘Start’ (chromatin start site): 0.0358, ‘5’UTR’: 0.0357, ‘gene_betweeness’ (gene length): 0.0267, ‘gene_degree’ (number of linked genes: 0.0249.Simultaneously, we calculated the contribution of different omics (somatic mutations, CNVs, RNA-seq data, DNA methylation) to identifying drug-available genes according to the Xgboost and random forest methods for evaluating the importance of multi-omics features (Fig. S9). The CNV-related features have the most considerable contribution to identifying cancer druggable genes, accounting for 55.6% and 59.5% in the Xgboost and random forest methods, followed by somatic mutations (43.3% calculated by the Xgboost and 35.4% calculated by the random forest). The contribution of RNA-seq and methylation features to identified cancer druggable genes is much smaller than that of CNVs and somatic mutations. Still, it is also helpful for druggable gene discovery.

**Supplementary Note 9 : The discussion**

The literature study revealed that a significant number of genes in the CDG-genes set obtained by DF-CAGE are supported by evidence. For instance, FoxA1 plays a crucial role in developing breast and prostate cancers. It is an attractive therapeutic target and has the potential to function as a novel biomarker. EP300 is also reported to serve as a prognostic marker and potential therapeutic target for TNBC. These publications provide ample evidence of the reliability of DF-CAGE.

Previous studies have shown that most cancer druggable genes are derived from cancer driver genes. In this study, the set of cancer druggable genes obtained by DF-CAGE and the known driver genes (CGC genes) have many intersections. We found that 215 of the 465 genes reported by DF-CAGE are also in the CGC set. However, some driver genes without evidence of being druggable, such as AFF4 and LCP1 (not in OncoKB, TARGET, DrugBank databases and not supported by drug-usable-related literature), were not reported by DF-CAGE. Hence, the advantage of DF-CAGE is the discovery of cancer druggable genes that are not cancer drivers (such as SMARCA1, ZNF311, ATXN3), which are also crucial for the development of targeted drugs.

By interpreting the DF-CAGE model results, we found that DF-CAGE focuses more on features obtained from CNVs while the driver gene discovery approaches mainly rely on somatic mutation-related features. We used the Random Forest and the XGboost for the model interpretation and yielded similar ranking results: 60% of the Top10 ranked features are related to copy number variations. We also found that genes closely associated with other genes are more likely to be druggable. Also, the analysis results revealed that the more the genes with a high proportion of mutations in the population, the more likely they are to be druggable, which may be because previous drug development took into account the broadest possible audience for the drug. The ability of the multi-omics data to identify cancer druggable genes with high accuracy reveals an inextricable link between cancer treatment and the multi-omics data.

The exploration of cancer driver genes is a significant bottleneck in cancer study, and some driver genes still exist that have not been identified. There are 194 genes in the DF-CAGE potential gene set, and although there is no evidence yet that these genes are druggable, some of these genes may be driver genes due to the close association between cancer druggable genes and driver genes. In contrast, of the 655 CGC cancer genes (often used as the gold standard for driver gene identification), 443 genes were not in the DF-CAGE prediction results, and 525 genes were not in the OncoKB, TARGET, or DrugBank cancer druggable genes sets, which suggests that only some of the driver genes may be druggable. Combining the above analysis, we consider a close association between driver genes and cancer druggable genes, but there is still a difference. With rapidly developing precision medicine, the exploration of the driver and cancer druggable genes will become more comprehensive.

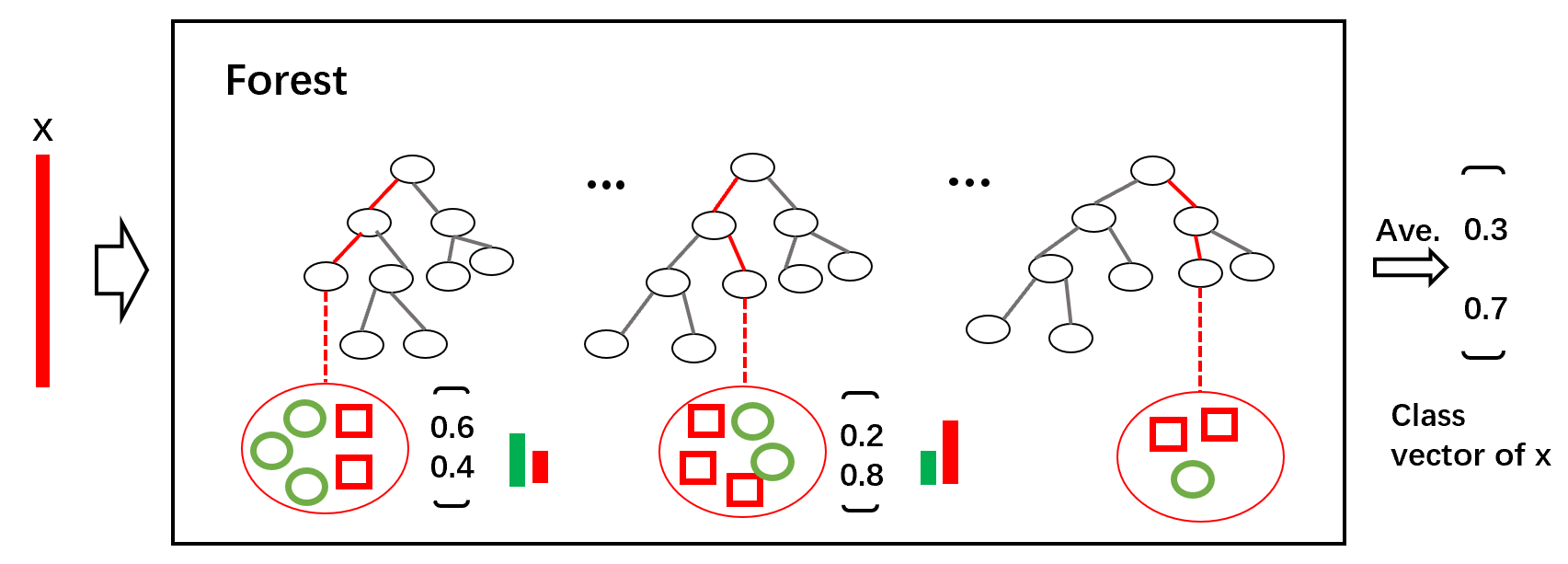

**Figure S1** The training process of DF-CAGE

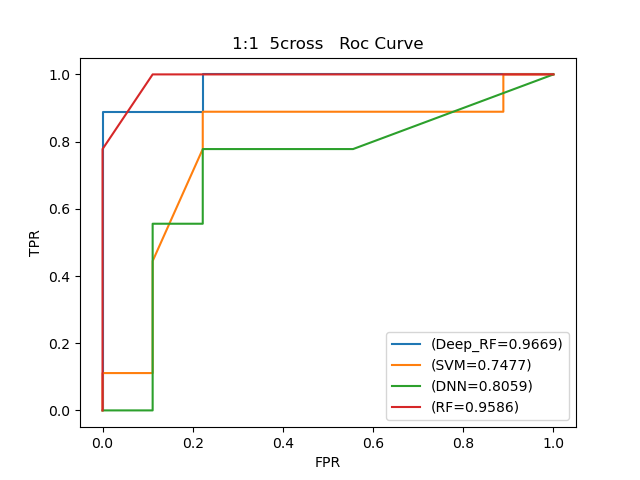

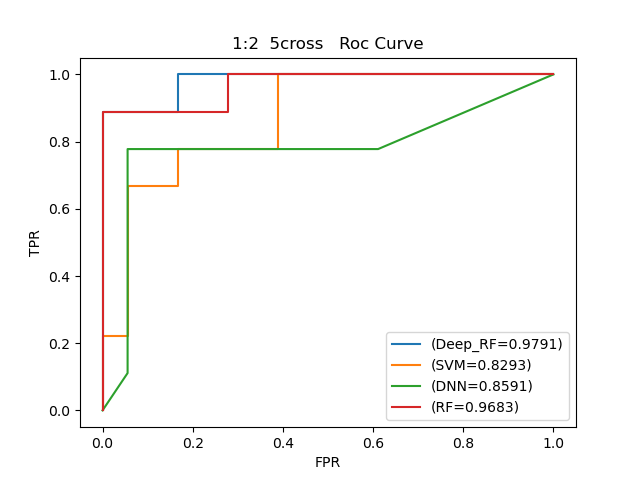

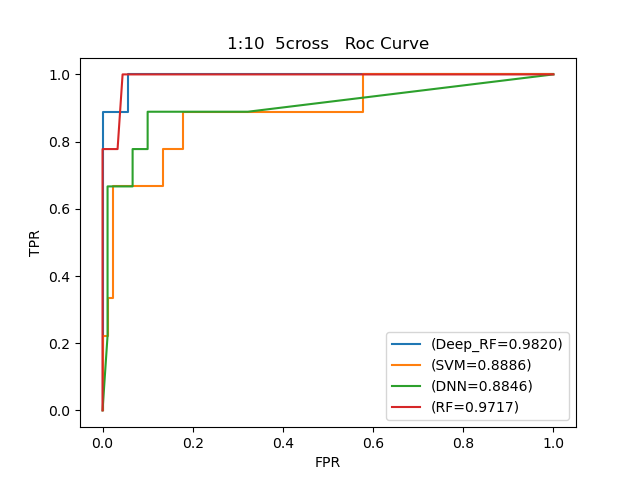

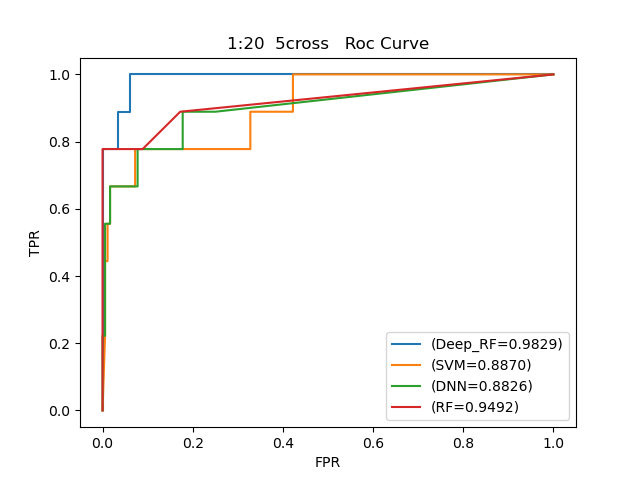

**Figure S2** Performance of different methods across the four test scenarios with various positive and negative sample ratios

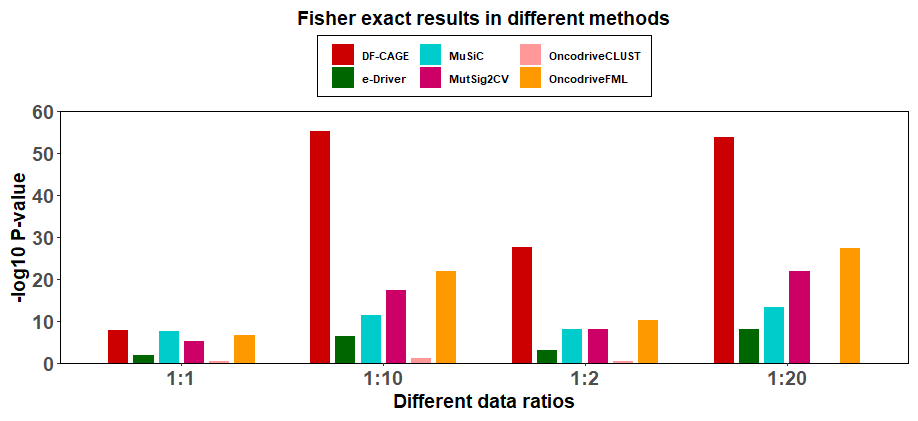

**Figure S3** The enrichment analysis of different methods across four test scenarios of the TARGET dataset

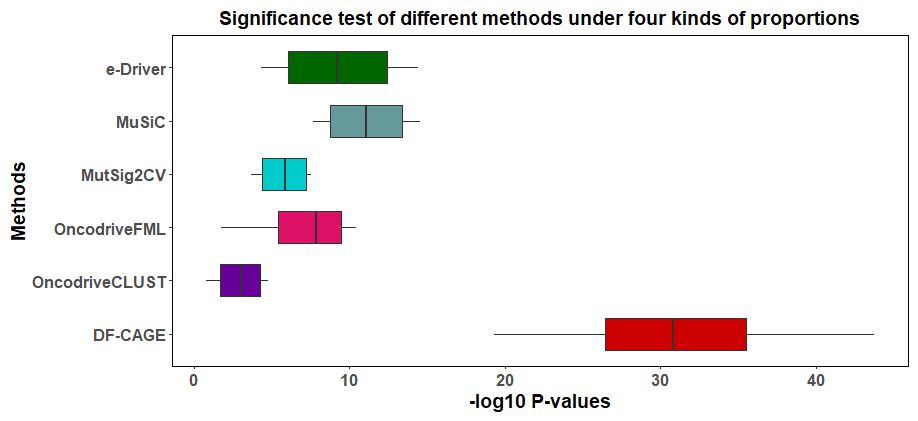

**Figure S4** The performance (Mann-Whitney test -log10 p-values) of different methods across four test scenarios of the TARGET dataset

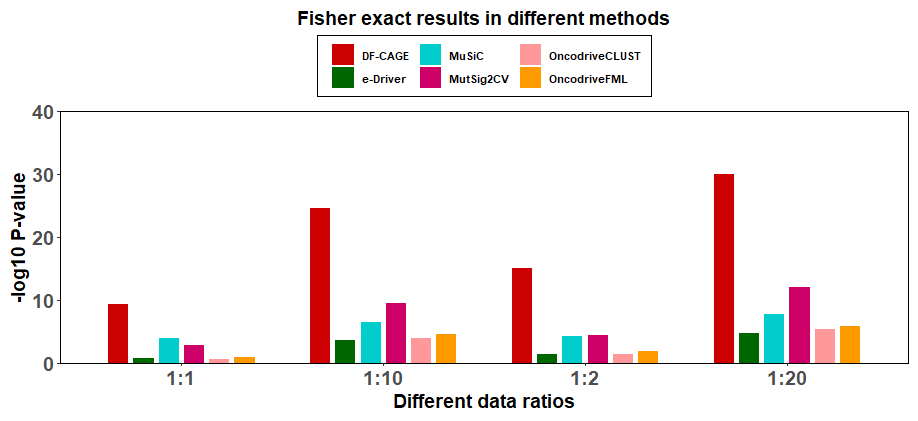

**Figure S5** The enrichment analysis of different methods across four test scenarios of the Drugbank dataset

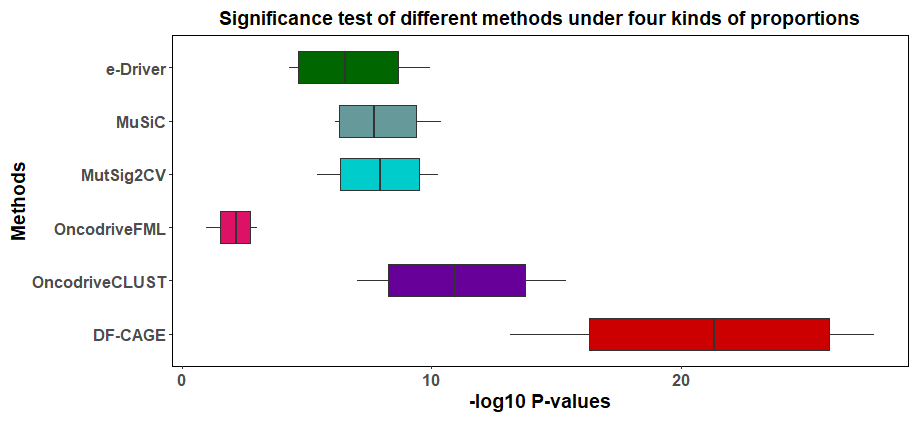

**Figure S6** The performance (Mann-Whitney test -log10 p-values) of different methods across four test scenarios of the Drugbank dataset

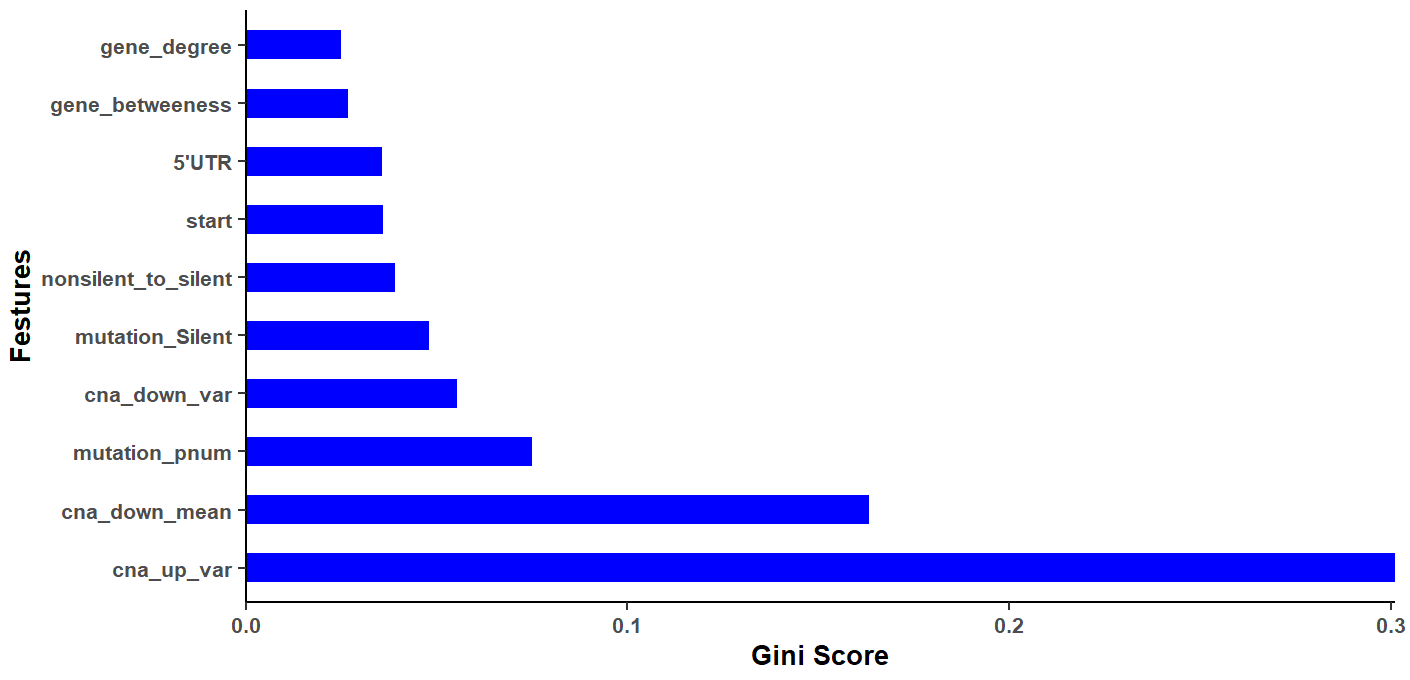

**Figure S7** Importance scores of DF-CAGE input features on the TARGET dataset

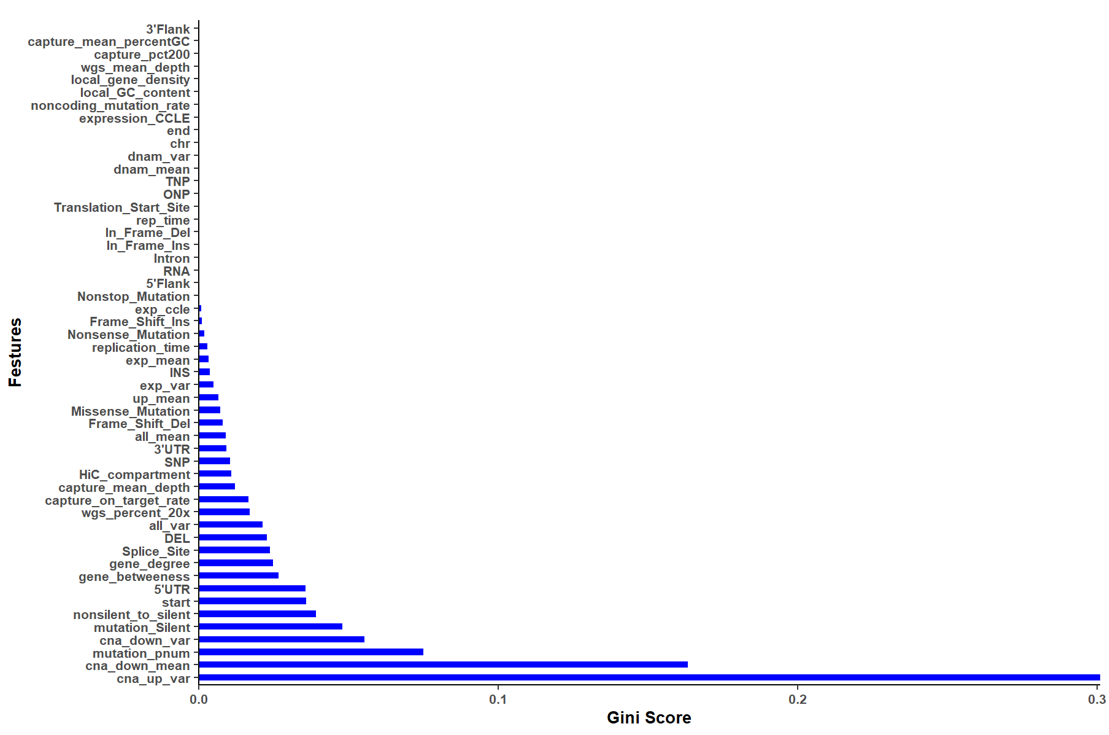

**Figure S8** Feature importance evaluation results for all the input of DF-CAGE

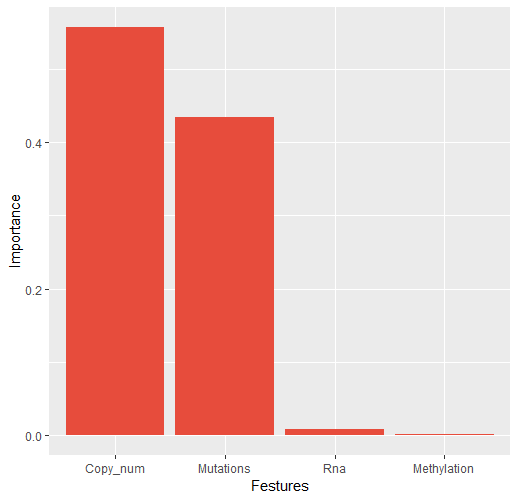

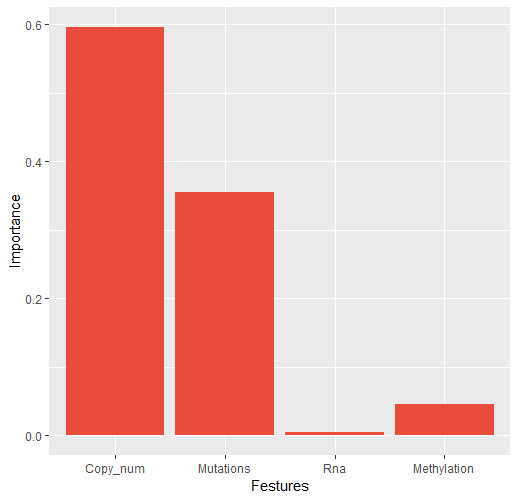

**Figure S9** Contributions of different omics data calculated by Xgboost (left) and random forest (right)

| **Methods** | **1:1** | **1: 2** | **1: 10** | **1: 20** |
| --- | --- | --- | --- | --- |
| **DF-CAGE** | **0.9669** | **0.9791** | **0.9820** | **0.9829** |
| **OncodriveCLUST** | **0.5565** | **0.6146** | **0.6568** | **0.7082** |
| **OncodriveFML** | **0.7398** | **0.7699** | **0.7729** | **0.7738** |
| **MutSig2CV** | **0.7827** | **0.8143** | **0.8174** | **0.8229** |
| **MuSiC** | **0.7842** | **0.8069** | **0.8613** | **0.8684** |
| **e-Driver** | **0.6291** | **0.6444** | **0.7307** | **0.7488** |

**Table S1** Performance (AUROC) of DF-CAGE and the comparison methods on the OncoKB dataset
